# Alcohol impacts an fMRI marker of neural inhibition in humans and rodents

**DOI:** 10.1101/2025.05.07.652668

**Authors:** Monami Nishio, Xinyi Wang, Eli J. Cornblath, Sung-Ho Lee, Yen-Yu Ian Shih, Nicola Palomero-Gallagher, Michael J. Arcaro, David M. Lydon-Staley, Allyson P. Mackey

## Abstract

Acute alcohol consumption leads to cognitive and behavioral disinhibition that increase health and social risks, such as traffic fatalities and violence. Although rodent studies have shown that alcohol affects inhibitory neuronal activity, its relevance to humans remains largely unexplored. Here, we used the Hurst exponent—an fMRI-based marker of neural inhibition—to examine alcohol’s effects in both rats and humans. In rats, acute alcohol administration significantly reduced the cortical Hurst exponent, suggesting a decrease in inhibitory neuronal activity. This reduction was strongly correlated with the spatial distribution of GABA_A_ receptor expression, highlighting the key role of these receptors in mediating alcohol’s effects. Similarly, in humans, acute alcohol consumption reduced the cortical Hurst exponent, especially in brain regions with high GABA_A_ receptor expression. Our findings provide cross-species *in vivo* evidence that acute alcohol consumption modulates neural inhibition, offering new insights into the neural mechanisms underlying alcohol-induced behavioral modulation.

## Introduction

Acute alcohol consumption triggers cognitive disinhibition, significantly increasing various health and social risks. For instance, approximately 37 people in the United States die daily in drunk-driving crashes, which account for 32% of all traffic fatalities ^1^. 40% of violent crimes, including assault, homicide, and domestic abuse, are committed by individuals with high blood alcohol content (BAC) at the time of their arrest ^2^. Moreover, nearly 30% of individuals who die by suicide consume alcohol shortly beforehand ^3^. In addition to causing cognitive impairment, acute alcohol intoxication disrupts motor coordination ^4,5^ and affects the brain’s reward systems ^6–9^, potentially leading to long-term issues such as binge drinking and addiction ^10^. Cognitive disinhibition caused by acute alcohol are often associated with weakened top-down control ^11^. However, the precise mechanism by which alcohol affects cortical neuronal activity, leading to disrupted top-down control, remains unclear.

*Ex vivo* studies using brain slices of rodents have shown that acute alcohol reduces the activity of excitatory neurons in the orbitofrontal ^12,13^, cingulate ^14^, and prefrontal ^15^ cortices. An *in vivo* study using multielectrode extracellular recordings also reported decreased spike activity ^15^, while a separate study using calcium imaging found increased activity in pyramidal neurons in the prefrontal cortex ^16^. The suppression of pyramidal neurons by alcohol is thought to involve both potentiation of GABA_A_ receptors and inhibition of glutamate receptors (NMDA, AMPA)^17,18^. While GABA_A_ receptor potentiation is a well-known effect of alcohol, the situation is more complicated, as certain GABA_A_ receptor subtypes are sensitive to alcohol at levels typically reached during social drinking, while other subtypes respond to higher concentrations associated with severe intoxication ^19^. Additionally, there is evidence that alcohol’s impact on GABA_A_ receptor activity is influenced by factors such as cell viability ^20^, cell type ^21^, and post-translational modifications like receptor phosphorylation ^22,23^. Furthermore, previous studies suggest that the suppression of pyramidal neurons is primarily attributed to its antagonistic effects on NMDA receptors, with minimal impact on AMPA receptor-mediated excitatory postsynaptic currents or GABA_A_ receptor-mediated inhibitory postsynaptic currents ^12,14^.

Fewer studies have examined alcohol’s effects on inhibitory neurons. One *ex vivo* study showed that alcohol reduces the activity of fast-spiking interneurons in the prefrontal cortex ^24^. Similarly, an *in vivo* calcium imaging study observed suppressed activity of somatostatin interneurons in the same region ^16^. Several *ex vivo* studies using rodent brain slices have demonstrated that the agonistic effect on GABA_A_ receptors is the primary mechanism through which alcohol affects inhibitory neurons ^25–27^.

In summary, rodent studies indicate that alcohol suppresses both excitatory and inhibitory neuronal activity, but through distinct mechanisms: suppression of excitatory neurons is mediated primarily by antagonistic effects on NMDA receptors, whereas suppression of inhibitory neurons is mediated primarily by agonistic effects on GABA_A_ receptors.

In humans, evidence of alcohol’s effects on neural activity is limited and indirect. Some studies using arterial spin labeling have shown that alcohol increases cerebral blood flow across widespread cortical regions ^28–31^. However, the relationship between neural activity and blood flow is complex and not fully understood ^32–34^.

Other research has found that alcohol consumption increases functional connectivity ^35–38^, especially within networks in the visual cortex ^36,39,40^, as well as in the sensorimotor cortex ^36^ and frontal cortex ^35^. Increased connectivity could result from various factors, including changes in neural activity—whether increased or decreased—or alcohol-induced vasoconstriction ^34,41,42^. Although the connection between these imaging findings and neural inhibition remains unclear, alcohol’s effects tend to be global rather than region-specific, impacting multiple cortical areas. Therefore, it is reasonable to expect alcohol to induce changes at the whole-brain level.

Understanding neural inhibition in humans requires appropriate non-invasive *in vivo* tools. Magnetic resonance spectroscopy can estimate GABA levels ^43^, but it can only measure one large voxel at a time, making it unsuitable for whole-brain analysis. An emerging alternative is the Hurst exponent ^44–46^, which quantifies long-range temporal correlations and scale-invariant dynamics in various time series ^47^. Computational models have demonstrated that enhancing inhibitory neuronal activities leads to an increase in the Hurst exponent of simulated local field potential and BOLD signals ^44–46,48^. Using LFP data from mice, Gao et al. showed that regions with a higher GABA synapse density have a higher Hurst exponent, indicating a spatial relationship between the inhibitory synaptic density and the Hurst exponent ^46^. Our recent study further revealed a strong spatial correlation between PV inhibitory neurons and the Hurst exponent across the cortex in both humans and rodents, suggesting a robust representation of inhibitory neurons across species ^49^. Given alcohol’s suppressive effects on inhibitory neurons observed in rodents ^16,24^, along with its widespread effects on the human brain reported in previous studies, we expect alcohol to reduce the Hurst exponent across brain regions.

Applicable across species, the Hurst exponent overcomes species-specific limitations. For instance, relying solely on human data may be confounded by factors like alcohol-induced motion artifacts or cognitive slowing. Additionally, individual responses to alcohol vary widely due to factors such as genetic differences ^50^, sex ^51^, lean body mass ^51^, liver volume ^51^, stress levels ^52^, and drinking history ^53^. In contrast, relying exclusively on rodent data presents challenges related to anesthesia and translating findings to humans. Previous acute alcohol studies have primarily focused on frontal regions in rodents, raising questions about their relevance to human brains and other cortical areas.

In this study, we first examine alcohol’s impact on the Hurst exponent in genetically and environmentally consistent, alcohol-naive, and age-controlled rodents to minimize individual variability. We also explore the correlation between alcohol’s effect on the Hurst exponent and GABA_A_ receptor expression across the cortex, shedding light on the receptors likely contributing to alcohol’s impact on neural inhibition. Next, we test the replicability of these findings in humans using an extensive repeated-measures (10 session) dataset. This study provides *in vivo* evidence of alcohol’s effect on neural inhibition across species.

## Results

### Alcohol Exposure Reduces the Hurst Exponent in Rats

We examined the effect of alcohol on the Hurst exponent in anesthetized rats. We parcellated the rat cortex into 50 parcels using the functional rat brain atlas ^54^. Using fractional integration models ^44,55,56^, we computed the autocorrelation function (ACF) of the time series for each cortical parcel, estimated the fractional integration order (*d*), and the Hurst exponent calculated as H = *d* + 0.5 (Figure 1A). Generally, neural signals and BOLD-fMRI data exhibit Hurst exponent values greater than 0.5, typically ranging between 0.6 and 0.9, indicating the presence of positive autocorrelation or persistent patterns in brain activity. A higher Hurst exponent, or a steeper slope, is believed to indicate greater neural inhibition^44–46^. A total of 38 Wistar rats underwent a single scanning session, during which 75 minutes of resting-state fMRI (rs-fMRI) BOLD data were collected (five 15-minute blocks)(Fig. 1B). After the first 15 minutes (baseline), each rat received a saline injection, followed by two 1 g/kg and one 2 g/kg injections of 20% ethanol, with each injection administered at the start of the subsequent 15-minute periods. This protocol resulted in cumulative ethanol doses of 0, 1, 2, and 4 g/kg over the course of the scan. To focus on steady-state data, we excluded the first 5 minutes of each block containing the injection periods, analyzing only the final 10 minutes of each block.

**Figure 1.**
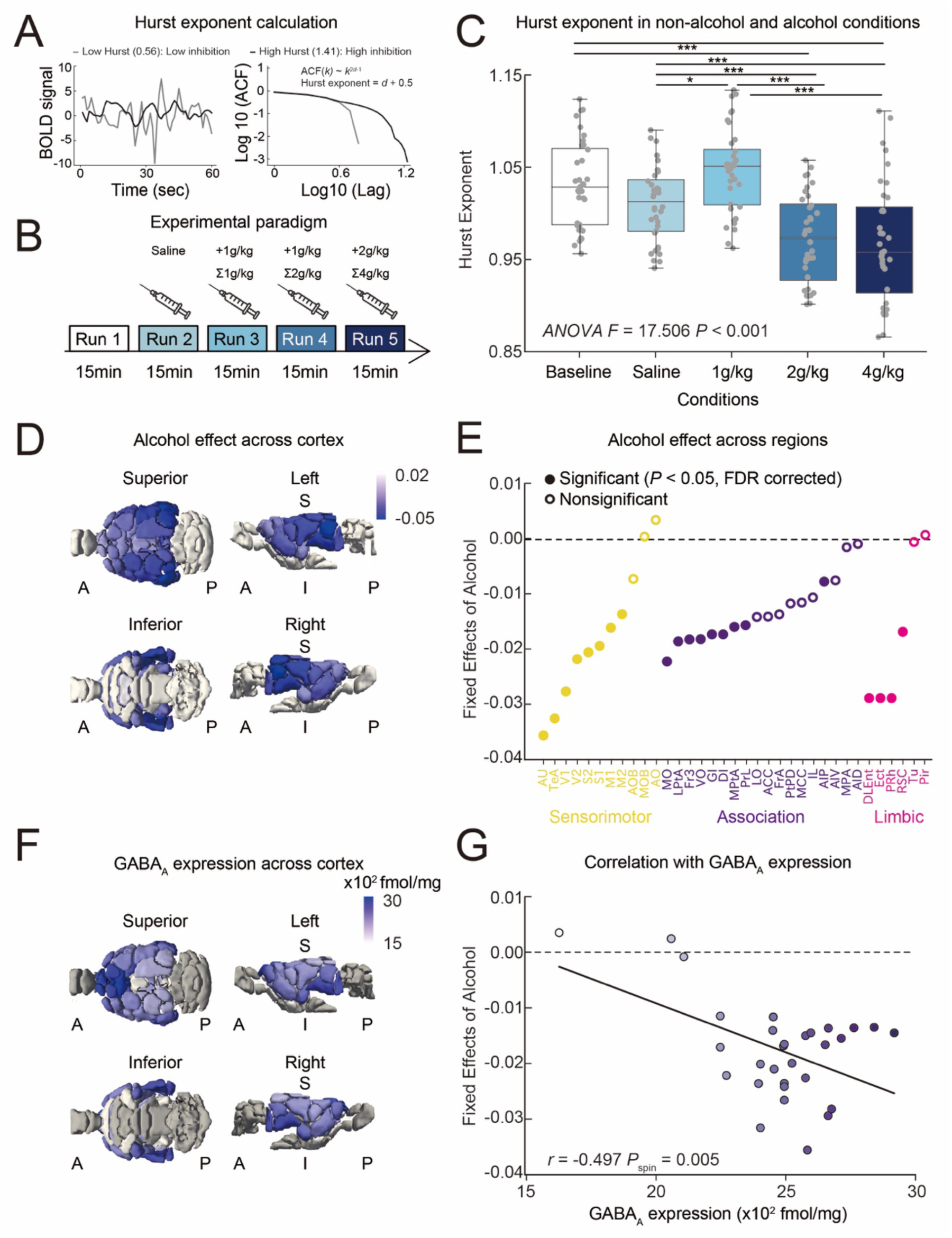
Alcohol’s effect on the Hurst exponent in rats. **(A)** Examples of fMRI time series with high and low Hurst exponents and autocorrelation function (ACF). ACF decays according to a power law, approximately as ACF(*k*) ∼ *k*^2d−1^, where *k* is the lag. H is defined as H = *d* + 0.5, with *d* representing the fractional integration order. **(B)** Schematic illustration of the experimental paradigm for rats. **(C)** The whole-brain averaged Hurst exponent in rats under different conditions (ANOVA *F* = 17.506, *P* < 0.001). To control for motion, we regressed out the FD mean from the original Hurst values and added the residuals to the average Hurst exponent. Post-hoc Tukey HSD test \*\*\**P* < 0.001, \*\**P* < 0.01, \**P* < 0.05. **(D)** Alcohol’s effect across the cortex in rats. **(E)** Alcohol’s effect in sensorimotor (yellow), association (purple) and limbic (pink) regions. Colored filled dots indicate regions with a significant alcohol effect that survived FDR correction, while unfilled dots represent regions with non-significant effects. The full names of the regions are listed in Table S1. **(F)** Cortical GABA_A_ receptor expression as visualized *in vitro* by means of the radioligand [^3^H] flumazenil. **(G)** Correlation between alcohol’s effect and GABA_A_ receptor expression across the cortex (*r* = −0.497, *P*_spin_ = 0.005).

Due to anesthesia, rat movement was generally minimal. However, three animals showed outlier levels of motion (mean framewise displacement exceeding 3 standard deviations above the mean) and were therefore excluded. After excluding these animals, there was a significant correlation between conditions (baseline, saline, and 1, 2, 4 g/kg alcohol) and mean framewise displacement (mean FD), suggesting that alcohol increases motion (Fig. S1A, ANOVA *F* = 4.576, *P* = 0.002). There was no significant correlation between the Hurst exponent and mean FD in any condition (Fig. S1B, *P*s > 0.069). After the 2 g/kg and 4 g/kg ethanol injections, the Hurst exponent was significantly lower compared to the first dose, saline, or baseline across the whole brain (Fig. 1C, ANOVA *F* = 17.506, *P* < 0.001). We examined region-specific effects across 50 cortical parcellations defined by the functional rat brain atlas ^54^ (Fig. 1D). The fixed effect of alcohol on the Hurst exponent was estimated using a linear mixed-effects model for each parcel, where the dose of alcohol (0, 1,2,4 g/kg), age, sex, and mean FD were treated as fixed effects. Random intercepts were modeled for each subject to account for repeated measures. The fixed effect of alcohol was extracted and its significance was calculated through an ANOVA test. The p-values was corrected for multiple comparisons using the false discovery rate (FDR) method. Since each brain region covers multiple parcels, the mean fixed effect per region is calculated across these parcels. A brain region was considered to have a significant effect if at least one parcel within the parcel survives FDR correction. The effect of alcohol remained significant after FDR correction with *P* < 0.05 in many brain regions across sensory, limbic, and association areas (Fig. 1E, the full names of the regions are listed in Table S1). The effect was particularly strong in auditory regions (auditory cortex, temporal association cortex)(Fig. 1E).

Since previous research suggests that alcohol’s effects on inhibitory neurons are primarily mediated through GABA_A_ receptors ^25–27^, we explored the correlation between alcohol’s effect on the Hurst exponent and GABA_A_ receptor expression across the cortex. To quantify GABA_A_ receptor density, we used *in vitro* visualization with flumazenil, the most commonly used ligand in PET studies of the GABAergic system ^57^. This analysis was conducted on rat brain data processed as part of a previously published study ^58,59^. Specifically, GABA_A_ receptor expression was determined in cortical regions defined by the atlas proposed by Haghir et al., 2024 ^60^. Out of the 50 regions of the functional rat brain atlas used in the fMRI study ^54^, 30 regions have available receptor density data (Fig. 1F). The Pearson correlation between alcohol effects and GABA_A_ receptor expression was then calculated for these 30 parcels. The significance of the correlation was assessed using spatially constrained rotation tests, known as ‘spin tests’ ^61^, because cortical data often exhibit distance-dependent spatial autocorrelation potentially inflating the significance of correlations between two cortical feature maps. There was a significant negative correlation between alcohol’s effect on the Hurst exponent and GABA_A_ receptor expression (Fig. 1G, *r* = −0.497, *P*_spin_ = 0.005), indicating that regions with higher GABA_A_ receptor expression experienced a greater reduction in the Hurst exponent. This result aligns with the previous findings that alcohol’s effect on inhibitory neurons is mainly mediated by GABA_A_ receptors ^25–27^.

Subcortical regions also exhibited a significantly lower Hurst exponent after 2 g/kg or 4 g/kg ethanol injections (Fig. S2A, ANOVA *F* = 9.109, *P* < 0.001), with the amygdala (central amygdala, basolateral amygdala), cerebellum, and striatum (dorsal, dorsomedial and ventrolateral parts of caudate putamen) surviving FDR correction with *P* < 0.05 (Fig. S2B).

### Alcohol Exposure Reduces the Hurst Exponent in Humans

Since rodent models offer valuable insights into neural mechanisms but do not fully capture the complexities of human brain function, we next investigated whether similar effects occur in humans. Eleven healthy adults (ages 24–33 years) participated in the experiment. Of these, eight completed 10 laboratory sessions—five with alcohol and five without (Fig. 2A). One participant completed an additional non-alcohol session, resulting in a total of 5 alcohol and 6 non-alcohol sessions. One participant completed only five scans (2 alcohol and 3 non-alcohol sessions), and another completed only four scans (2 alcohol and 2 non-alcohol sessions). During alcohol sessions, participants consumed alcoholic beverages over a 30-minute period to reach a blood alcohol content (BAC) of 0.08%—roughly equivalent to three standard drinks—and were then taken directly to the MRI scanner. In the non-alcohol sessions, participants did not consume alcohol prior to the MRI. The Hurst exponent was calculated from 7-minute runs of rs-fMRI data for both conditions. Since alcohol increased participant movement (Fig. S3A, *t* = −3.677, *P* < 0.001), particularly at moderate BAC levels (Fig. S3B), and the Hurst exponent significantly correlated with motion (Fig. S3C, Non-alcohol; *r* = −0.811, *P* < 0.001, Alcohol; *r* = −0.814, *P* < 0.001), we excluded sessions with a mean framewise displacement (mean FD) greater than 0.3 mm (Fig. S3C) and controlled for mean FD for subsequent analysis. We examined region-specific effects across 100 cortical parcellations defined by the Schaefer 100 atlas ^62^. The fixed effect of alcohol on the Hurst exponent was estimated using a linear mixed-effects (LME) model for each parcel, where alcohol condition (alcohol / non-alcohol session), person-average mean FD, run-specific variation from the person-average mean FD, and gender were treated as a fixed effects, while a random slope and intercept for alcohol was be included for each subject to model individual variability in the effect of alcohol on the hurst exponent.

**Figure 2.**
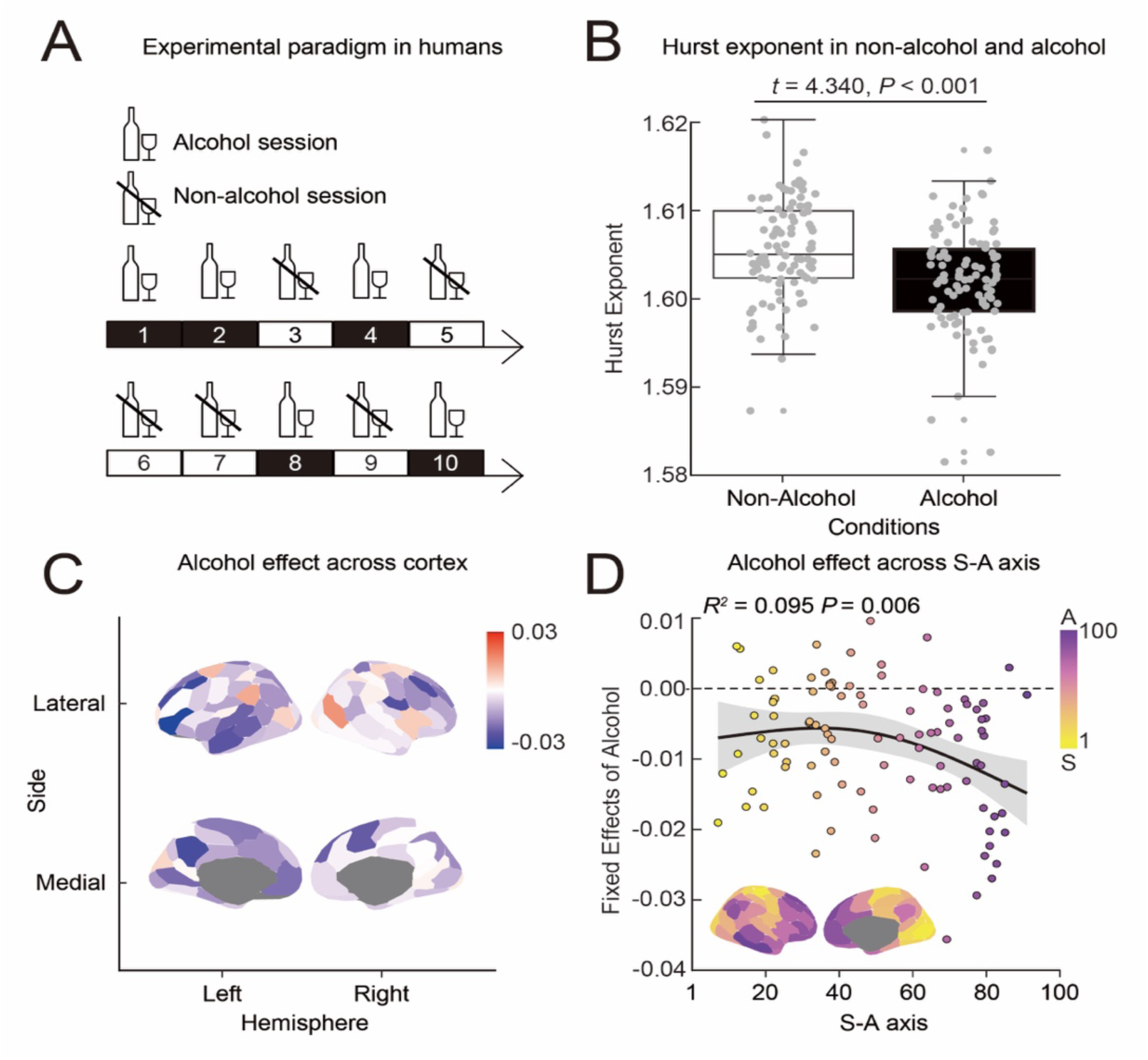
Alcohol’s effect on the Hurst exponent in humans. **(A)** Schematic illustration of the experimental paradigms for humans. Healthy adults (ages 24–33) participated in 10 laboratory sessions—five with alcohol and five without. The order of alcohol and non-alcohol sessions was randomized. **(B)** The whole-brain averaged Hurst exponent in humans under non-alcohol (white) and alcohol (black) conditions (*t* = 4.340, *P* < 0.001). To control for motion, we regressed out the FD mean from the original Hurst values and added the residuals to the average Hurst exponent. **(C)** Alcohol’s effect across the cortex. **(D)** Mapping of alcohol’s effect along the sensory (yellow)-association (purple) axis (*R*^2^ = 0.095, *P* = 0.006).

Consistent with rodent findings, the average cortical Hurst exponent was significantly lower during alcohol sessions compared to non-alcohol sessions (Fig. 2B, *t* = 4.340, *P* < 0.001). This reduction remained significant using stricter motion exclusion criteria (mean FD < 0.2) (Fig. S3D, E, *t* = 3.140, *P* = 0.002). To assess regional specificity, we tested alcohol’s effect on 100 evenly sized cortical parcels (28). Although no parcels survived FDR correction, the effect was consistently negative across the cortex (Fig. 2C). The most pronounced decreases were observed in association regions (Fig. 2D, GAM *R*^2^ = 0.095, *P* = 0.006). This pattern remained even when non-alcohol sessions were subsampled to balance session counts between alcohol and non-alcohol sessions (Fig. S3F, *R*^2^=0.063, *P*=0.029).

For the correlation analysis between alcohol’s effects and receptor expression, we used the dataset published by Hansen et al., 2022 ^63^. Volumetric PET images were collected from multiple studies for 19 different neurotransmitter receptors, including GABA_A_ receptors. These images provide an estimate proportional to receptor densities, so the measured values (such as binding potential and tracer distribution volume) are simply referred to as receptor density. All PET images were registered to the MNI-ICBM 152 non-linear 2009 (version c, asymmetric) template and parcellated into 100 parcels according to the Schaefer 100 atlas ^62^.

GABA_A_ receptors were expressed throughout the cortex, with particularly strong expression at both ends of the S-A axis (Fig. 3A, B, *R²* = 0.144, *P* < 0.001). GABA_A_ receptor expression significantly negatively correlated with alcohol’s effect on the Hurst exponent, suggesting that regions with higher GABA_A_ receptor expression experienced a greater reduction in the Hurst exponent due to alcohol. Since both alcohol’s effects and GABA_A_ receptor expression exhibit a significant correlation with the S-A axis (Fig. 2E, 3B), correlation between alcohol’s effects and GABA_A_ receptor expression could be partly explained by their independent associations with the S-A axis. However, the correlation between alcohol’s effect and GABA_A_ receptor expression remained significant even after controlling for the S-A axis with partial correlation test (Fig. 3C *r* = −0.320, *P* = 0.001), indicating that the S-A axis does not fully explain this relationship. Similarly, we examined the correlation between alcohol’s effect and the expression of various receptors, uptake sites and transporters while controlling for S-A axis. Among all, GABA_A_ ranked the highest (Fig. 3D). Serotonergic receptors (5HT2A *r* = −0.297, *P* = 0.003, 5HT6 *r* = −0.246, *P* = 0.014, 5HT1b *r* = −0.243, *P* = 0.016, 5HTT *r* = −0.218, *P* = 0.030), dopamine transporter (DAT *r* = −0.275, *P* = 0.006), and metabotropic glutamate receptor 5 (mGluR5 *r* = −0.255, *P* = 0.011) also showed significant correlations with alcohol’s effect.

**Figure 3.**
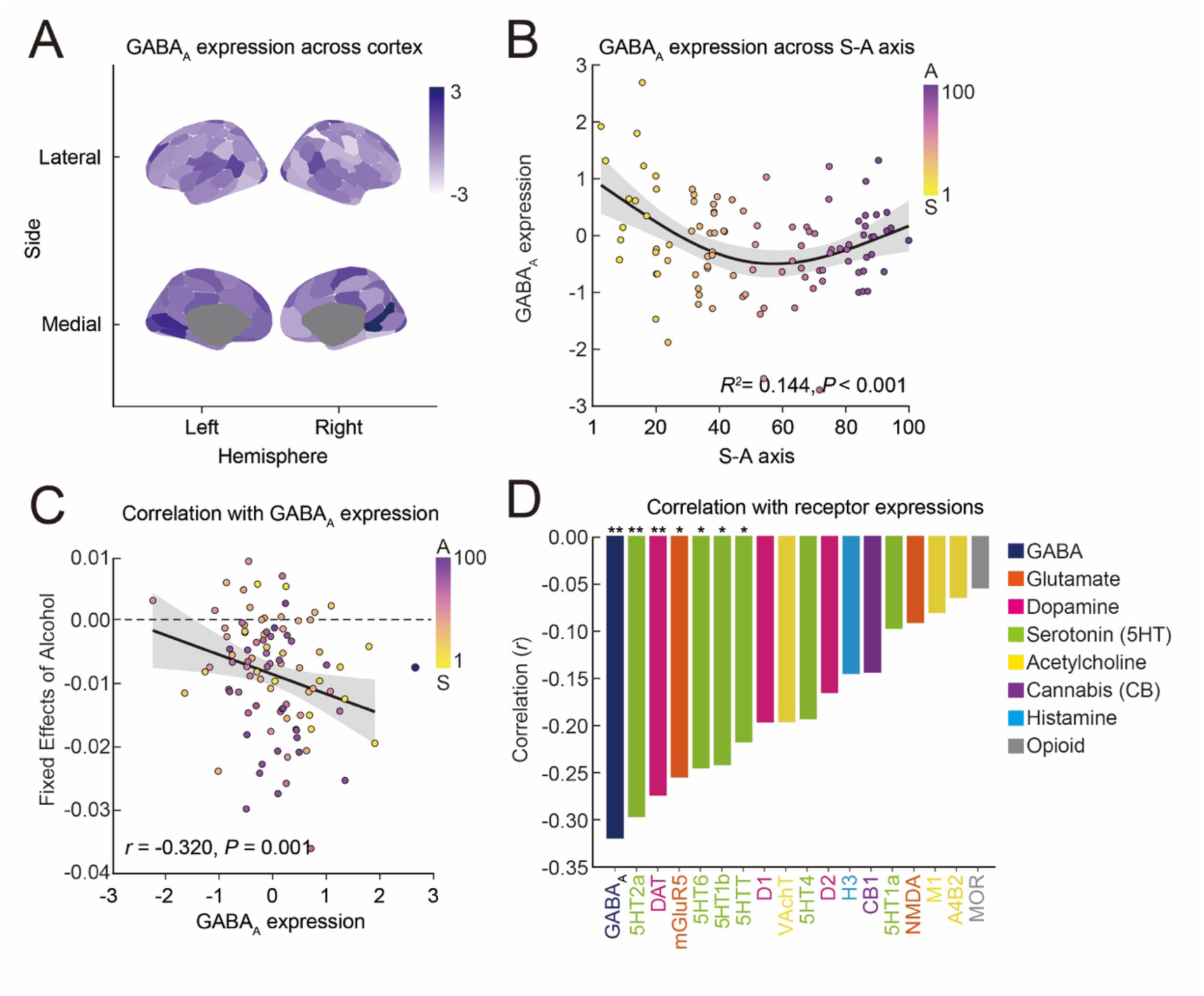
Alcohol’s effect on the Hurst exponent in humans. **(A)** GABA_A_ receptor expression across the cortex (z-scored). **(B)** Mapping of GABA_A_ receptor expression along the sensory (yellow)-association (purple) axis (*R²* = 0.144, *P* < 0.001). **(C)** Correlation between alcohol’s effect and GABA_A_ receptor expression across the cortex. S-A rank is controlled using partial correlation test (*r* = −0.320, *P* = 0.001). **(D)** Comparison of correlation coefficients between group-averaged alcohol effect and different type of receptors. S-A rank is controlled using partial correlation test. \**P* < 0.05, \*\**P* < 0.01. The color of the bar indicates the type of receptor. Dark blue: GABA, Orange: Glutamate, Pink: Dopamine, Green: Serotonin (5HT), Yellow: Acetylcholine, Purple: Cannabis (CB), Light blue: Histamine, Gray: Opioid.

Alcohol also significantly reduced the Hurst exponent in subcortical regions (Fig. S2C, *t* = 10.36, *P* < 0.001). Although no parcels survived FDR correction, the effect was most pronounced in the cerebellum, globus pallidus, nucleus accumbens, and substantia nigra (Fig. S2D).

## Discussion

In this study, we investigated the effects of alcohol on the Hurst exponent—an fMRI-based marker of neural inhibition—in both rats and humans. Our findings revealed a consistent reduction in the Hurst exponent across species, indicating that alcohol decreases inhibitory neuronal activity. Moreover, the spatial distribution of alcohol’s effects was strongly correlated with GABA_A_ receptor expression, consistent with a role for GABA_A_ receptors in alcohol’s effects.

Most prior studies on alcohol’s effects on cortical neuronal activities have focused on prefrontal regions in rodents ^12–16,24^, leaving its impact on other brain areas largely unexplored. Here, we demonstrated that alcohol’s effects are widespread across the cortex. Studies using arterial spin labeling have similarly reported increased cerebral blood flow across broad cortical regions ^28–31^, suggesting alcohol’s widespread impact on the cortex. Additionally, alcohol has been shown to enhance intrinsic connectivity within networks in the visual cortex ^36,39,40^, as well as in the sensorimotor cortex ^36^ and frontal cortex ^35^. Given the consistent reduction in the Hurst exponent within these sensorimotor and association areas, we speculate that decreased inhibition may lead to the disinhibition of excitatory neurons, increasing their spontaneous activity and potentially enhancing within-network connectivity. Interestingly, there are subtle differences in the spatial patterns of alcohol effects across species. In rats, the alcohol effect was strongest in sensory regions, while in humans, it was strongest in association regions. This could be due to differences in brain organization between species, with sensory regions being more dominant in rats and association regions playing a larger role in humans.. This finding may, to some extent, align with the global reduction in brain volume observed in humans, even among light drinkers (only one to two alcohol units per day), with the most pronounced effects in association regions ^64^. Although the relationship between acute changes in neuronal activities and structural alterations has not been studied, one possible explanation is that neuronal hyperexcitability could lead to excitotoxicity, resulting in neuronal death and, ultimately, structural volume reduction.

Consistent with previous studies identifying GABA_A_ receptors as key mediators of alcohol’s effects on inhibitory neurons ^25–27^, we found a significant spatial correlation between alcohol’s effects and GABA_A_ receptor expression in both rodents and humans. However, there was considerable variability in the relationship between GABA_A_ receptor expression and alcohol effects in rats, as certain frontal regions—including the anterior cingulate cortex, frontal association areas, prelimbic cortex, primary motor cortex (M1) and premotor cortex (M2)— exhibited high GABA_A_ receptor expression but only moderate reductions in the Hurst exponent following alcohol exposure. This discrepancy may stem from variations in GABA_A_ receptor subtypes, which have differing sensitivities to alcohol ^19–23^. Additionally, in humans, we found that the serotonin 2A receptor (5HT2A) and dopamine transporter (DAT) were strongly correlated with alcohol’s effects (*P* < 0.01). Several studies have confirmed an ethanol dose-dependent increase in dopamine release in both rodents and humans ^6,8,9,65–68^. Regions with high DAT expression, indicating dense dopamine innervation, are likely to be most affected by alcohol-induced increases in dopamine. Since dopamine dose dependently suppresses inhibitory neuronal activity through D2 receptors ^69–71^, these regions may experience a pronounced impact of alcohol on inhibitory neuronal suppression. Alcohol is also known to interact with serotonin receptors ^72–74^. While the effects of alcohol on excitatory neurons are primarily mediated via NMDA receptors ^12,14^, we found no correlation between alcohol’s effects and NMDA receptor expression. Instead, we observed a significant correlation between alcohol’s effects and mGluR5. Both NMDA receptors and mGluR5 are closely localized on the cell surface, with mGluR5 activation enhancing NMDA receptor activity ^75–78^. Therefore, it is possible that alcohol’s effect on the Hurst exponent could be influenced by its impact on excitatory neurons through mGluR5, which may, in turn, modulate NMDA receptor activity.

We also observed a significant reduction in the Hurst exponent across several subcortical regions in rats, including basal ganglia (CPuD, CPuM, CPuVL) and cerebellum. The reduction in inhibition and overactivation of Purkinje cells due to the blockade of adenosine uptake in the cerebellum is known to cause motor dysfunction, referred to as ataxia, which appears immediately after acute alcohol consumption ^5^. The alcohol-induced increase in striatal dopamine release (50–56) is also possibly due to the suppression of inhibitory neurons and the subsequent overactivation of dopamine neurons ^7^. Although the effects did not reach statistical significance, the basal ganglia and cerebellum also show the largest alcohol-related effects among subcortical regions in humans. However, the validity of the Hurst exponent as a marker of neural inhibition in subcortical regions remains uncertain, as no studies have yet applied this metric to these areas. Additionally, the ROIs for subcortical regions varied significantly in size compared to the cortical parcellations. Therefore, comparing the Hurst exponent across these regions should be done with caution.(45–47).

Our study has several limitations. One challenge in using the Hurst exponent is its susceptibility to motion artifacts, particularly in conditions where motion differs significantly between conditions, such as in alcohol studies. Compared to rats, alcohol’s effects on the Hurst exponent in humans were relatively subtle, as no individual cortical or subcortical regions showed significant effects after correction for multiple comparisons. This may be due to the strong correlation between motion and the Hurst exponent, as alcohol increases motion (presumable by reducing behavioral inhibition), and controlling for motion may have overly diminished the observed alcohol effect. Well-documented alcohol-induced changes in the cardiovascular system both in rodents and humans ^79–81^ could also contribute to increased motion, underscoring the need for better understanding and control of these factors. In rats, isoflurane anesthesia was used, which potentiates GABA_A_ receptor function. Although anesthesia conditions were consistent between alcohol and non-alcohol sessions, and our results aligned with findings from awake human participants, it remains possible that anesthesia interacted with alcohol’s effects in rodents. Replicating our findings in awake rodents or under different anesthesia types would strengthen these conclusions. Furthermore, some animal studies suggest that alcohol’s effects on inhibitory neuronal activity are dose-dependent ^16^ Consistent with this, we observed dose-dependent effects on the Hurst exponent in rats. Specifically, 1 g/kg did not cause a significant change, whereas 2 g/kg and 4 g/kg significantly reduced the Hurst exponent. Our study used a blood alcohol concentent of 0.08% in humans, which is the legal driving limit and considered relatively high. Our findings may differ at lower doses; therefore, future research should investigate these dose-dependent effects in humans. Finally, to enhance statistical power in our group-level analysis with a relatively small sample size, we used a repeated-measures design. While this approach is appropriate for estimating main effects of alcohol, the sample size was not large enough for investigating between-individual variability, such as differences related to gender, age, or genetic background.

In summary, this study provides *in vivo* evidence of alcohol’s suppressive effects on neural inhibition across species. Given that most alcohol studies are conducted in rodents, the use of consistent methods across species and the consistency of our cross-species findings may be critical for future translational research ^82^. These insights could be valuable in informing clinical interventions and preventive strategies to address the health risks associated with alcohol consumption.

## Methods and Materials

## 1. Rat data and analyses

### 1.1 In vivo Imaging subjects

The data analyzed in this study were previously published by Lee et al. ^37^. Briefly, a total of 38 Wistar rats, bred and reared in-house, were included in the experiment. The pups remained with their mother until weaning at postnatal day 21 (P21), and were subsequently scanned at either postnatal day 45 (P45; N = 19, 11 females) or postnatal day 80 (P80; N = 19, 10 females). All animals were maintained in a temperature- and humidity-controlled vivarium under a 12:12 h light-dark cycle, with ad libitum access to food and water.

### 1.2 Experimental design

Each animal underwent a single scanning session at its designated age (P45 or P80). During each session, a total of 75 minutes of blood oxygenation level-dependent (BOLD) resting-state fMRI (rs-fMRI) data were collected, divided into five 15-minute blocks. After the first 15 minutes of baseline scanning, animals received an intraperitoneal injection of saline, followed by two 1 g/kg and one 2 g/kg injections of 20% v/v ethanol, with one injection administered per 15-minute block ^10,83^. Rats were initially anesthetized with 4% isoflurane, which was then reduced to 2% for secure placement in an MR-compatible rat cradle. Once stabilized, a continuous infusion of dexmedetomidine (0.05 mg/kg/hr) and pancuronium bromide (0.5 mg/kg/hr) was administered, while isoflurane was further reduced to 0.5%. This infusion protocol was initiated 30 minutes prior to the start of rs-fMRI scans ^84^.

### 1.3 MRI data acquisition

All MR images were acquired at the UNC Center for Animal MRI (CAMRI) on a Bruker BioSpec 9.4-T, 30 cm bore system using an 86 mm volume coil (RF transmitter) and 4-channel receiver. Magnetic field homogeneity was optimized with global shimming followed by local second-order MAPSHIM. BOLD rs-fMRI data were collected with a 2D multi-slice, single-shot, gradient-echo EPI sequence (TR = 2000 ms, TE = 14 ms, flip angle = 70°, FOV = 28.8 × 28.8 mm, matrix = 72 × 72, 32 slices, 0.4 mm isotropic voxels).

### 1.4 MRI data preprocessing

All rs-fMRI data were first corrected for slice timing and motion using AFNI ^85^. Time-averaged images were used to generate brain masks via an in-house 2D-UNET algorithm ^86^ and manually refined with ITK-SNAP ^87^. These masks were applied to the corresponding rs-fMRI data to remove non-brain signals, followed by spatial normalization to an in-house EPI template using ANTs SyN registration ^88^. Nuisance signals, including baseline trends and six motion parameters, were regressed out, and the data were band-pass filtered (0.01–0.1 Hz) and smoothed with a 0.5 mm FWHM Gaussian kernel. Each session was divided into five 15-minute blocks corresponding to different exposure periods. Data were parcellated using a functional rat brain atlas ^54^, and time courses were averaged within parcels to calculate the Hurst exponent. Correspondence between parcels and anatomical regions is provided in Supplementary Table 3 of Lee et al., 2021^54^.

### 1.5 Hurst exponent

The Hurst exponent can be estimated using several methods. The Power Spectral Density (PSD) method leverages the relationship between the spectral density of the form 1/f*^β^* and the Hurst exponent *α*, where *β* = 2*α*−1 ^89^. Alternatively, Detrended Fluctuation Analysis (DFA) estimates the Hurst exponent by quantifying how fluctuations in a detrended, integrated time series scale with the size of the observation window, providing robustness against nonstationarities commonly present in neural data ^90–94^. Finally, fractional integration models treat the time series as fractionally integrated processes, where the Hurst exponent H is defined as H = *d* + 0.5, with *d* representing the fractional integration order ^44,55,56^. In these models, the autocorrelation function (ACF) decays according to a power law, approximately as ACF(*k*) ∼ *k*^2d−1^, where *k* is the lag. This means that for *d* > 0, the correlations between distant observations decline slowly, reflecting long-range dependence or persistence in the signal. This approach fits parametric models to capture long-range dependencies more rigorously. For this study, we employed the fractional integration method in our analysis to ensure robust and stable estimation. In this study, Hurst exponent was computed using the *nonfractal* MATLAB toolbox (https://github.com/wonsang/nonfractal)^44,55^. The specific function utilized is bfn_mfin_ml.m function with the ‘filter’ argument set to ‘haar’ and the ‘ub’ and ‘lb’ arguments set to [1.5,10] and [−0.5,0], respectively. This toolbox uses a discrete wavelet transform and a model of the time series as fractionally integrated processes (FIP) and is estimated using maximum likelihood estimation. FIP model was employed instead of the Fractional Gaussian Noise (fGn) model to eliminate the assumption of stationarity and the upper bound of H = 1.

### 1.6 Alcohol effect modeling

To model the effect of alcohol, we employed a linear mixed-effects (LME) approach similar to that used in humans. For each brain region in the dataset, we constructed a model using the formula ‘Regional Hurst exponent ∼ Run + Age + Sex + mean FD + (1 | Subject)’, where the run (saline, dose1, dose2, dose3), age, sex, and mean FD were treated as fixed effects. We selected the saline condition as the control instead of the baseline condition to eliminate the influence of the injection procedure on the Hurst exponent. Random intercepts were modeled for each subject to account for repeated measures. To calculate the alcohol effect for each brain region, we grouped the parcels belonging to each brain region and computed the average Hurst exponent of those parcels.

### 1.7 Receptor expression

For the correlation analysis between alcohol effects and receptor expression, we used receptor autoradiographs generated in the framework of a study published by Palomero-Gallagher et al. ^58,59^. Eight adult male LEW/Ztm rats were included in the study, with serial 20 µm coronal sections obtained at Bregma levels 4.20, 1.56, −2.76, and −4.80 ^95^. Sections were alternately stained for cell bodies using a modified silver Nissl-staining method or processed for quantitative *in vitro* autoradiography for receptor visualization. Receptor labeling involved pre-incubation in a Tris-citrate buffer to remove endogenous substances, incubation with [^3^H]-flumazenil to identify total binding sites, and a rinsing step to stop the binding process. Flumazenil reveals the GABA_A_-associated benzodiazepine (GABA_A_/BZ) binding sites and is the ligand used the GABA_A_ receptor in the receptor PET dataset published by Hansen et al. ^63^. Sections were exposed to beta-radiation-sensitive films alongside calibrated microscales to enable densitometric analysis of the ensuing receptor autoradiographs. The autoradiographs were digitized, their grey values converted into receptor densities (fmol/mg protein) by means of the microscales, and pseudo-color coded for visualization. Regions of interest (ROIs) were identified using the adjacent Nissl-stained sections, Bregma levels from the Paxinos and Watson rat brain atlas, and cortical parcellation by Haghir et al ^60^ to ensure precise boundaries, and manually annotated on the color-coded images. Receptor densities were extracted with AnaRec software^96^, which extracted the receptor density measures from each ROI for further analysis.

For each parcel included in the functional rat brain atlas ^54^, we aggregated the GABA_A_ receptor expression from the regions within the parcel and calculated their average. Consequently, receptor density data were available for 30 out of 50 parcels. We then computed the Pearson correlation between the alcohol effects and GABA_A_ receptor expression for these 30 parcels.

## 2. Human data and analyses

### 2.1 Imaging participants

Participants meeting fMRI inclusion and study inclusion criteria (N = 11; M = 28.1 years, SD = 3.26 years; 4 women, 7 men; 3 identified as Asian, 5 identified as White, 1 identified as White and Hispanic or Latinx) were recruited. The study inclusion criteria was over the age of 21, have consumed at least 1 beverage containing alcohol during the 6 months before their first scan and have no physical conditions that could be influenced by drinking alcohol. The study’s exclusion criteria included women who were pregnant, planning a pregnancy, or lactating. Individuals with a history of cancer, heart disease, stroke, or myocardial infarction within the past six months were also excluded. Participants with contraindications against fMRI, a history of problematic alcohol use (defined as an AUDIT score of 8 or higher), or liver disease were not eligible. Additionally, individuals taking medications for which alcohol use is contraindicated, as outlined by the National Institute on Alcohol Abuse and Alcoholism (NIAAA), were excluded from the study^97^. All participants provided written informed consent. Procedures were approved by the University of Pennsylvania Institutional Review Board.

### 2.2 Experimental design

Participants attended one baseline survey visit and 10 laboratory visits where each session included surveys and an MRI session. For 5 of the visits, participants were asked not to consume alcohol 12 hours before the visit and not to eat food for 3 hours prior to session. They were then provided with alcoholic beverages designed to increase their blood alcohol content (BAC) to 0.08% before they underwent MRI. The beverages consisted of vodka and orange juice mixed in a 3:1 juice:alcohol ratio. The amount consumed was equivalent to approximately 3 standard drinks of alcohol, though it was slightly more or less depending on the participant’s age, gender, height, and weight as calculated through a BAC calculator (https://dionysus.psych.wisc.edu/open/bac_calc.html). Participants consumed the beverage within 30 minutes and were immediately scanned, with BAC measured by breathalyzer before and after the MRI. For the other 5 visits, participants were not provided with alcoholic beverages prior to the MRI scan. Among the 11 participants, one completed an additional non-alcohol session, resulting in a total of 5 alcohol and 6 non-alcohol sessions. One participant completed only five scans (2 alcohol and 3 non-alcohol sessions), and another completed only four scans (2 alcohol and 2 non-alcohol sessions). The two participants who did not complete all 10 sessions left the study because their eligibility for participating in the study changed (n = 1) and because they could no longer make time for the study (n = 1). The ordering of alcohol and non-alcohol sessions was randomized for each participant.

### 2.3 MRI data acquisition

Imaging data were acquired at University of Pennsylvania on a 3T Siemens Magnetom Prisma scanner equipped with a 32-channel head coil. High-resolution T1-weighted structural images were obtained using a multi-echo magnetization-prepared rapid gradient-echo (MEMP-RAGE) sequence (repetition time (TR) = 2530 ms, echo time (TE) = 1.69/3.55/5.41/7.27 ms, TI = 1330 ms, FA (degree) = 7°, field of view (FoV) = 256 mm^2^, voxel-size =1 mm^3^, slice thickness = 1 mm, echo spacing = 11.1 ms, bandwidth = 650 Hz/pixel, and acquisition time = 5.26 min). During the structural scan, participants watched a 7-minute video featuring abstract shapes and moves in slow continuous transitions^98^ in order to help reduce movement, boredom and nervousness. For rs-fMRI, 60 transversal slices were acquired by using an echoplanar imaging sequence (TR = 800 ms, TE = 30 ms, FA (degree) = 52°, field of view (FoV) = 216 mm^2^, voxel-size = 2.4 mm^3^, slice thickness = 2.4 mm, echo spacing = 0.51mm, bandwidth =2778 Hz/pixel, and acquisition time = 7.04 min). Each session included four resting-state scans with 2 fixation resting-state scans and 2 film clip resting-state scans. For this study, we only used the fixation resting-state scans. During these scans, a white fixation cross was presented on a black screen.

### 2.4 MRI data preprocessing

The T1-weighted (T1w) image underwent intensity nonuniformity correction using N4BiasFieldCorrection^99^ from the Advanced Normalization ToolS (ANTS) version 2.2.0, and used as T1w reference throughout the workflow. Following this, skull stripping was performed using the antsBrainExtraction.sh script (ANTS version 2.2.0), with the OASIS template as the target. The brain mask was refined with a custom variation of the method to reconcile ANTS-derived and FreeSurfer ^100^-derived segmentations of the cortical gray matter of Mindboggle (RRID:SCR_002438^101^). Spatial normalization to the ICBM 152 Nonlinear atlases version 2009c. (RRID:SCR_008796^102^) was performed through nonlinear registration with antsRegistration (ANTS, version 2.2.0, RRID:SCR_004757^103^, using brain-extracted versions of both T1w volume and template. Brain tissue segmentation of CSF, white matter (WM), and gray matter was performed on the brain-extracted T1w using fast (Functional MRI of the Brain Software Library (FSL) version 5.0.9; RRID:SCR_002823^104^).

For each rs-fMRI run, a reference volume and its skull-stripped version were generated using a custom methodology of fMRIPrep version 20.2.0 (RRID:SCR_016216^105^), which is based on Nipype version 1.1.7 (RRID:SCR_002502^106^, nipy/nipype:1.1.7). Brain surfaces were reconstructed using the recon-all command ^100^ before other processing, and reconstructed surfaces were used as input to fMRIprep. The BOLD reference was then coregistered to the T1w reference using bbregister (FreeSurfer), employing boundary-based registration^107^. Coregistration was configured with nine degrees of freedom to account for distortions remaining in the BOLD reference. Head-motion parameters with respect to the BOLD reference (transformation matrices and six corresponding rotation and translation parameters) were estimated before any spatiotemporal filtering using the mcflirt tool (FSL version 5.0.9^108^, and slice-time correction was performed using 3dTshift from AFNI 20160207 (RRID:SCR_005927^109^). The BOLD time series were resampled onto the MNIPediatricAsym standard space by applying a single, composite transform, generating a preprocessed BOLD run in MNI152NLin6Asym space.

Several confounding time series were calculated based on the preprocessed BOLD, including FD, DVARS (root-mean-square intensity difference from one volume to the next), and three region-wise global signals (CSF, WM, and the whole brain). FD and DVARS were calculated for each functional run, both using their implementations in Nipype ^110^. The head-motion estimates calculated in the correction step were also placed within the corresponding confounds file.

All resamplings can be performed with a single interpolation step by composing all the pertinent transformations (i.e., head-motion transform matrices and coregistrations to anatomic and template spaces). Gridded (volumetric) resamplings were performed using antsApplyTransforms (ANTS), configured with Lanczos interpolation to minimize the smoothing effects of other kernels ^111^.

Further preprocessing was performed using a confound regression procedure that has been optimized to reduce the influence of participant motion ^112,113^; preprocessing was implemented in XCP-D 0.5.0 ^112^, a multimodal tool kit that deploys processing instruments from frequently used software libraries, including FSL^108^ and AFNI^85^. Further documentation is available at https://xcp-d.readthedocs.io/en/latest/ and https://github.com/PennLINC/xcp_d. Using XCP-D, the preprocessed functional time series on the fsLR cortical surface underwent nuisance regression, using a 36-parameter model that included three global signals (whole brain, CSF, and WM) and six motion parameters, as well as their derivatives, quadratic terms, and derivatives of quadratic terms. Linear regression, implemented in Scikit-Learn (0.24.2), was employed to regress the confounds. Motion censoring was applied, with outlier volumes exceeding FD = 0.3 mm flagged and removed from confound regression. An interpolated version of the BOLD data is created by filling in the high-motion outlier volumes with cubic spline interpolated data, as implemented in *nilearn*. Any outlier volumes at the beginning or end of the run are replaced with the closest non-outlier volume’s data, in order to avoid extrapolation by the interpolation function. If there was less than 100 seconds of data remaining after removing high-motion outlier volumes, those samples are excluded. No bandpass filtering was performed.

### 2.5 Hurst exponent

We used the same metrics as those applied to humans to calculate the Hurst exponent for rats.

### 2.6 Alcohol effect modeling

To model the effect of alcohol on the Hurst exponent, we employed a linear mixed-effects (LME) approach to account for the nested nature of the data (i.e., several runs nested in 11 participants) using the ‘lmer’ function from the ‘lme4’ package in R. For each brain region in the dataset, we constructed a model with the form ‘Regional Hurst exponent ∼ Alcohol + psm_mean_fd + wps_mean_fd + Gender + (1 + Alcohol | Subject)’, where alcohol condition (alcohol / non-alcohol session), psm_mean_fd (person-average mean FD), wps_mean_fd (run-specific variation from the person-average mean FD), and gender were treated as fixed effects, while a random slope and intercept for alcohol were included for each subject to account for individual variability in the response to alcohol. The optimization was performed using the ‘optimx’ method with the “nlminb” algorithm. For each region, we extracted the fixed effect of alcohol and calculated its significance through an ANOVA test. The p-values were then corrected for multiple comparisons using the false discovery rate (FDR) method.

### 2.7 Alignment with the S-A axis

We used the S-A axis derived by Sydnor and colleagues (31). This map encompasses various cortical hierarchies, including functional connectivity gradients, evolutionary cortical expansion patterns, anatomical ratios, allometric scaling, brain metabolism measures, perfusion indices, gene expression patterns, primary modes of brain function, cytoarchitectural similarity gradients, and cortical thickness.

### 2.8 Correlation with receptor expression

To calculate the correlation between alcohol’s effect and GABA_A_ receptor expression across the cortex, we computed the Pearson correlation across 100 cortical parcels. We compared the correlation coefficients and p-values for the GABA_A_ receptor and alcohol effect with those of other types of receptors to assess the specificity of the GABA_A_ receptor in explaining the spatial distribution of alcohol’s effect.

### 2.9 Spin-based spatial permutation testing

To address distance-dependent spatial autocorrelation in cortical data, we evaluated the significance of Pearson’s correlations between two whole-brain cortical feature maps (the Hurst exponent) using non-parametric spin tests. These tests, also known as spatially constrained rotation tests, compare the observed correlation to a null distribution obtained by spatially iterating (spinning) one of the feature maps. Specifically, the null distribution is generated by rotating spherical projections of one map while preserving its spatial covariance structure. The *P* value (*P*_spin_) is determined by comparing the empirical correlation to the distribution obtained from 10,000 spatial rotations. Spin tests were conducted using the spin permutation test algorithm available in the ENIGMA toolbox for Python (https://enigma-toolbox.readthedocs.io/en/latest/pages/08.spintest/index.html).

### 2.10 Receptor expression

For the correlation analysis between alcohol effects and receptor expression, we used the dataset published by Hansen et al. ^63^. Volumetric PET images were collected for 19 different neurotransmitter receptors and transporters across multiple studies. These images provide an estimate proportional to receptor densities, so we refer to the measured values (such as binding potential and tracer distribution volume) simply as receptor density. All PET images were registered to the MNI-ICBM 152 non-linear 2009 (version c, asymmetric) template and parcellated into 100 regions according to the Schaefer atlas.

## Acknowledgements

We thank all of the individuals who participated in this research. This study was supported by a National Science Foundation CAREER award (to A.P.M. under Grant No. 2045095), Brain & Behavior Research Foundation (to D.M.L.-S.), Quad Fellowship (to M.N.) and Nakajima Foundation Scholarship (to M.N.).

---

**Supplementary Figure 1.**
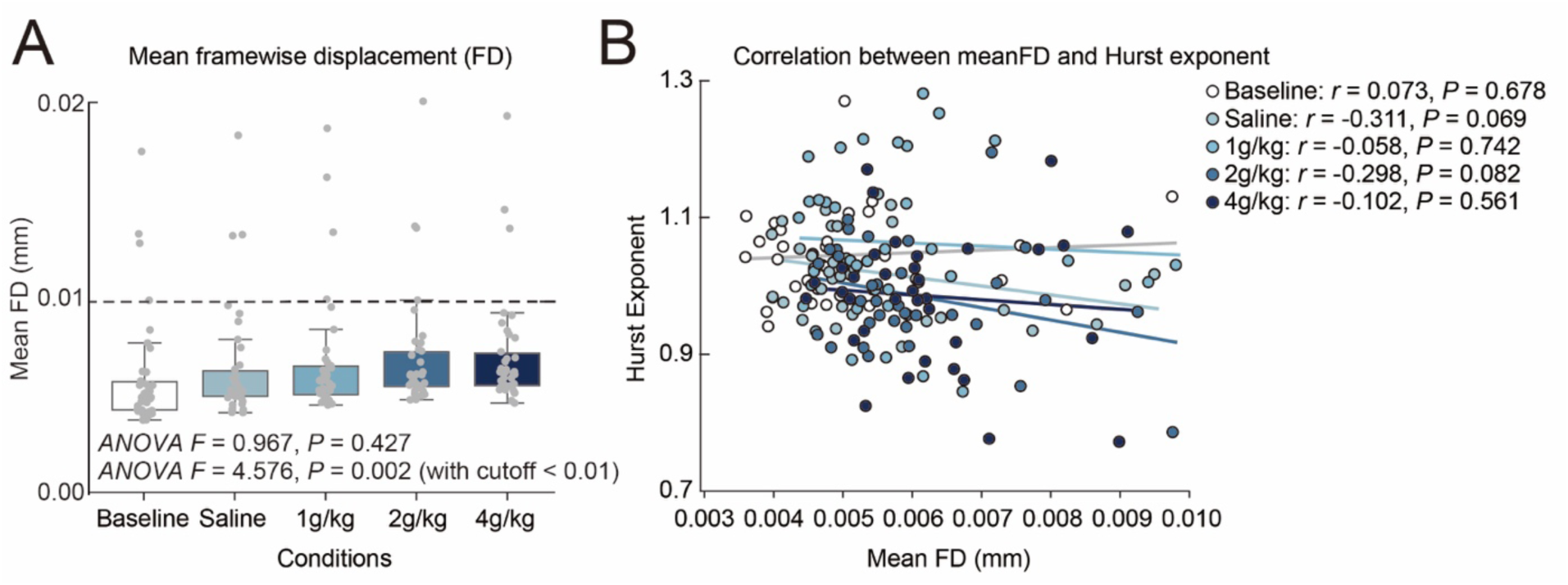
Correlation between the Hurst exponent and motion in rats. **(A)** Mean framewise displacement (mean FD) for each condition in rats without a cutoff (ANOVA *F* = 0.967, *P* = 0.427) and with a cutoff of 0.01 mm (ANOVA *F* = 4.576, *P* = 0.002). The dashed line indicates a cutoff of 0.01 mm. **(B)** Correlation between mean FD and the Hurst exponent after applying the motion cutoff (0.1 mm). Baseline (white): *r* = −0.063, *P* = 0.709, Saline (cyan): *r* = −0.075, *P* = 0.653, 1 g/kg (light blue): *r* = −0.023, *P* = 0.890, 2 g/kg (blue): *r* = - 0.043, *P* = 0.800, 4 g/kg (dark blue): *r* = −0.002, *P* = 0.991).

**Supplementary Figure 2.**
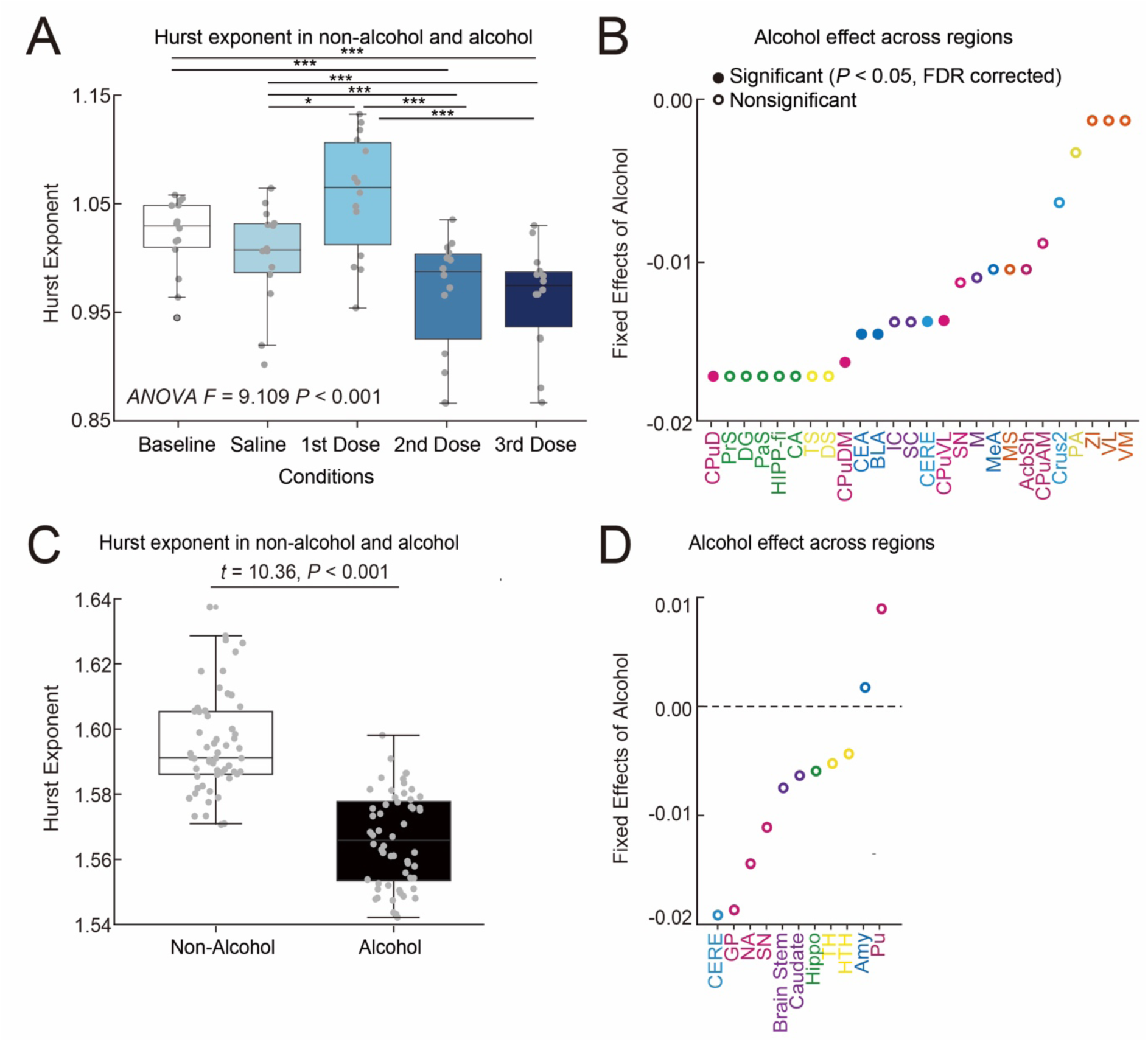
Alcohol’s effect on the Hurst exponent of subcortical regions. **(A)** The Hurst exponent averaged across subcortical regions in rats (ANOVA *F* = 9.109, *P* < 0.001). To control for motion, we regressed out the FD mean from the original Hurst values and added the residuals to the average Hurst exponent. Post-hoc Tukey HSD test \*\*\**P* < 0.001, \*\**P* < 0.01, \**P* < 0.05. **(B)** Alcohol’s effect across subcortical regions. Colored filled dots indicate regions with a significant alcohol effect that survived FDR correction, while unfilled dots represent regions with non-significant effects. Pink: Striatum, Green: Hippocampal, Yellow: Thalamic, Dark blue: Amygdala, Purple: Midbrain, Light blue: Cerebellum, Orange: Zonality. The full names of the regions are listed in Table S1. **(C)** The Hurst exponent averaged across subcortical regions in humans under non-alcohol (white) and alcohol (black) conditions (*t* = 10.36, *P* < 0.001). To control for motion, we regressed out the FD mean from the original Hurst values and added the residuals to the average Hurst exponent. **(D)** Alcohol’s effect across subcortical regions in humans. Unfilled dots represent regions with non-significant effects. Pink: Striatum, Green: Hippocampal, Yellow: Thalamic, Dark blue: Amygdala, Purple: Midbrain, Light blue: Cerebellum. The full names of the regions are listed in Table S1.

**Supplementary Figure 3.**
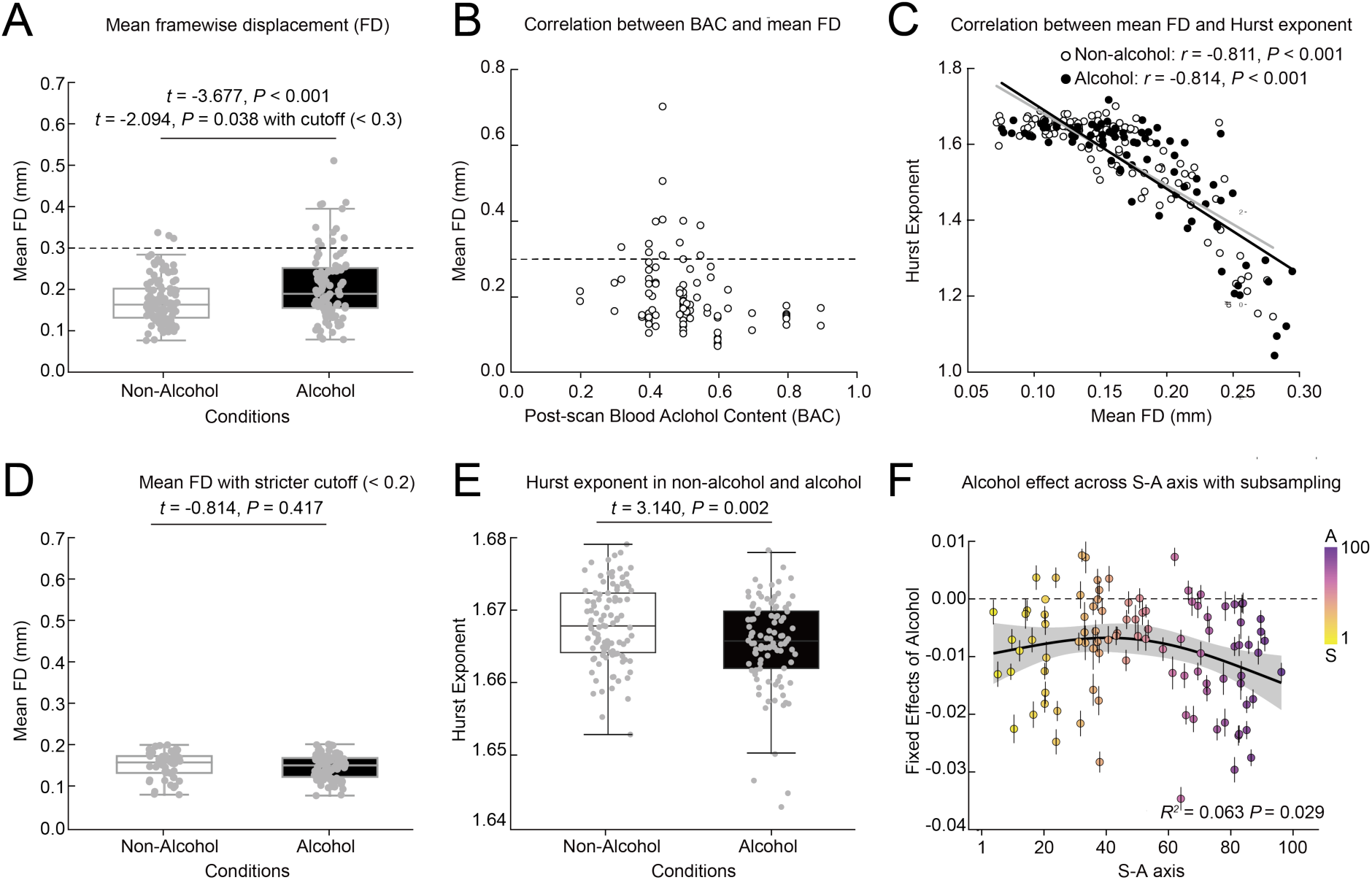
Correlation between the Hurst exponent and motion in humans. **(A)** Mean framewise displacement (mean FD) in non-alcohol and alcohol conditions without a cutoff (*t* = −3.677, *P* < 0.001) and with a cutoff of 0.3 mm (*t* = −2.094, *P* = 0.038). The dashed line indicates a cutoff of 0.3 mm. **(B)** Correlation between post-scan blood alcohol content (BAC) and mean FD. The dashed line indicates a cutoff of 0.3 mm. **(C)** Correlation between mean FD and the Hurst exponent after apply the motion cutoff (0.3 mm). White dots represent non-alcohol sessions, and black dots represent alcohol sessions (Non-alcohol: *r* = −0.811, *P* < 0.001, Alcohol: *r* = −0.814, *P* < 0.001). **(D)** Mean framewise displacement (mean FD) in non-alcohol and alcohol conditions with a stricter cutoff of mean FD < 0.2 mm (*t* = −0.814, *P* = 0.417). **(E)** The whole-brain averaged Hurst exponent with a stricter cutoff (*t* = 3.140, *P* = 0.002). **(F)** Alcohol’s effect across the cortex with balancing session counts between alcohol and non-alcohol conditions by subsampling the non-alcohol sessions (*R*^2^ = 0.063, *P* = 0.029). The error bar represents the standard deviation across 10 subsamples.

**Supplementary Table 1.**
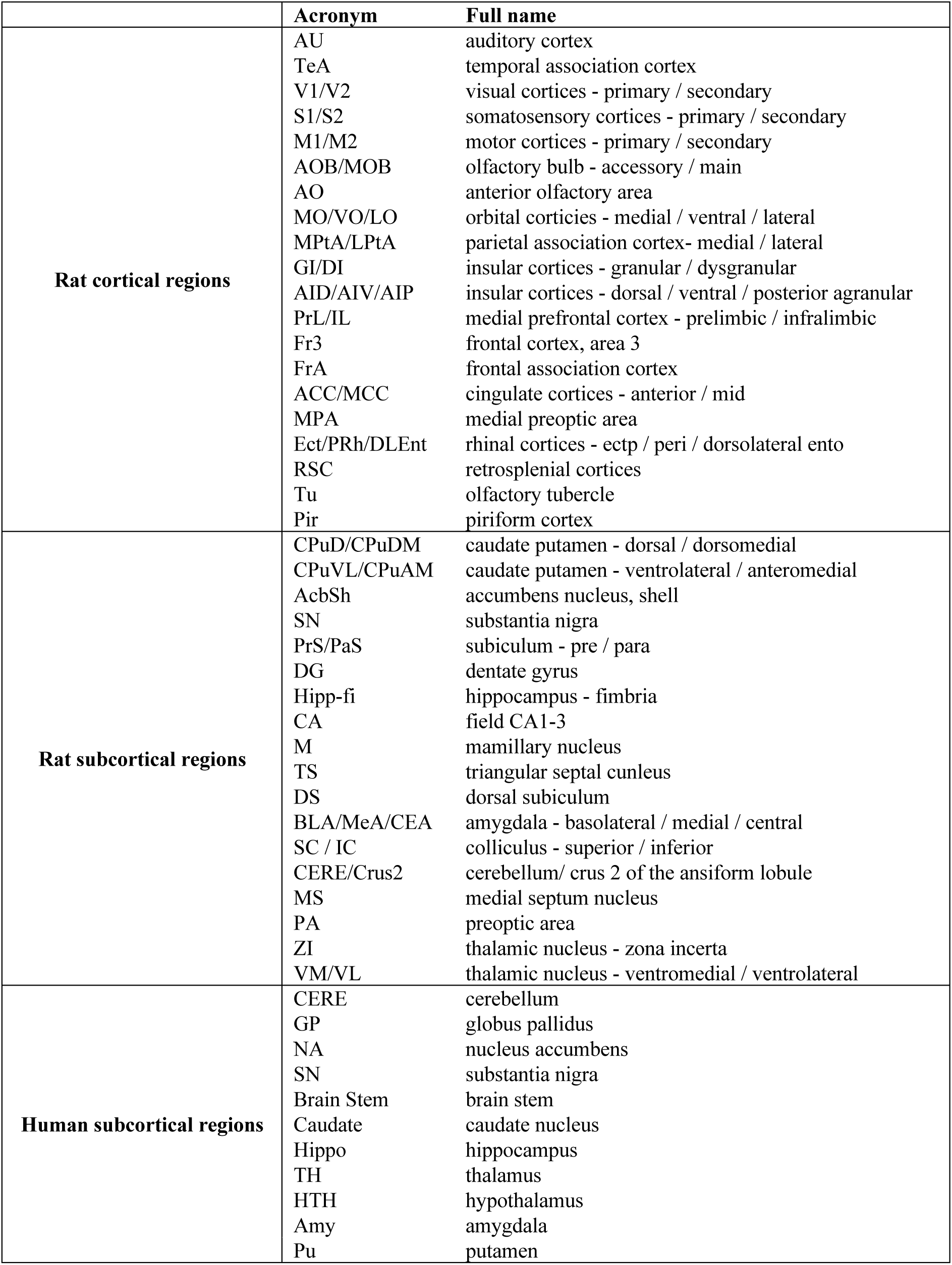
Full names of the brain regions.

